# A Bayesian model for genomic prediction using metabolic networks

**DOI:** 10.1101/2023.03.12.532311

**Authors:** Akio Onogi

**Affiliations:** Department of Life Sciences, Faculty of Agriculture, Ryukoku University, Otsu, Shiga, 520-2194, Japan

## Abstract

**Motivation:** Genomic prediction is now an essential technique in breeding and medicine, and it is interesting to see how omics data can be used to improve prediction accuracy. Precedent work proposed a metabolic network-based method in biomass prediction of Arabidopsis; however, the method consists of multiple steps that possibly degrade prediction accuracy

**Results:** We proposed a Bayesian model that integrates all steps and jointly infers all fluxes of reactions related to biomass production. The proposed model showed higher accuracies than methods compared both in simulated and real data. The findings support the previous excellent idea that metabolic network information can be used for prediction.

**Availability and implementation:** All R and stan scripts to reproduce the results of this study are available at https://github.com/Onogi/MetabolicModeling.

**Supplementary information:** This study provides no supplementary information

## 1. Introduction

Genomic prediction is a statistical method for predicting phenotypes or genetic merits using genome-wide DNA markers (Meuwissen *et al*., 2001). Genomic prediction is now essential in animal breeding (e.g., Garcia-Ruiz *et al*., 2016; Scott *et al*., 2021), and is becoming increasingly important in plant breeding (e.g., Dreisigacker *et al*., 2021). In breeding, selection strategies employing genomic prediction are often referred to as genomic selection. The framework of genomic prediction is also used in medicine to predict disease risks, which are known as polygenic or genetic risk scores (Wray *et al*., 2019; Lello *et al*., 2020).

One of the primary goals of genomic prediction research has been to improve prediction accuracy (e.g., Heslot *et al*., 2012; Onogi *et al*., 2015; Dreisigacker *et al*., 2021). This interest has motivated various statistical methods and models, including Bayesian regressions (reviewed in de los Campos *et al*., 2013; Dreisigacker *et al*., 2021). It has also been explored how to utilize information other than the genome to enhance prediction accuracy. The typical additional information is “omics data.” For example, transcriptome (Li *et al*., 2019; Perez *et al*., 2022) and metabolome (Riedelsheimer *et al*., 2012; Campbell *et al*., 2021) are often used together with the genome, and some studies have used both (Xu *et al*., 2016; Schrag *et al*., 2018). Furthermore, because data in biology are essentially multivariate (i.e., multiple traits can be measured for a genotype at multiple environments), it has been of interest to learn how to utilize multivariate information to improve prediction accuracy for target traits (e.g., Jia and Jannink, 2012; Jarquin *et al*., 2016; Atanda *et al*., 2022).

Upon such trends, Tong *et al*., (2020) proposed utilizing metabolic networks to predict the biomass of *Arabidopsis thaliana* (Arabidopsis). The method is based on the concept of flux balance: which states that the fluxes (amounts) of each reaction related to biomass production are determined in such a way that the production and consumption of each metabolite are balanced. Under this constraint, the flux of biomass production is estimated using quadratic programming such that it matches the observed biomass. When predicting the biomass of a new genotype, the fluxes of all reactions and biomass production are first predicted using ordinal genomic prediction methods, and the predicted fluxes are then adjusted using quadratic programming to fill the flux balance.

Although their findings suggest that metabolic network information can be used to predict phenotypes, their approaches still have room for improvement. First, because the flux is estimated for each genotype independently, dependencies (or covariances/similarities) between genotypes are not used. Second, because flux estimation and prediction are done separately, uncertainty in estimation increases noise during prediction. To overcome these drawbacks, the current study proposes a Bayesian model that estimates fluxes of all genotypes simultaneously and conducts flux estimation and prediction jointly. The performance of the proposed model was compared with other methods including the approach of Tong *et al*., (2020). The results demonstrated the superiority of the proposed model and supported the excellent idea presented by Tong *et al*., (2020) that metabolic network information can aid in phenotype prediction.

## 2. Systems and methods

Let **M**_*i*_ denote the stoichiometry matrix of genotype *i* (Fig. 1A). **M**_*i*_ is a *K* by *J* matrix containing reaction coefficients, where *K* and *J* denote the number of metabolites and reactions, respectively. Let the *J*_th_ reaction be the one related to the target trait, and the other reactions (reaction *j* where 1 ≤ *j* < *J*) represent processes that produce metabolites related to the target trait. Here the target trait is the biomass of Arabidopsis. Let *V_i,j_* denote the flux of reaction *j* of genotype *i* which represents the amount that the reaction occurs in the genotype’s cells/tissues and let **V** be an *N* by *J* matrix containing all fluxes of all genotypes, where *N* is the number of genotypes. **V**_*i*_ is the *i*_th_ rows of **V** which denotes the fluxes of genotype *i*.

**Fig. 1.**
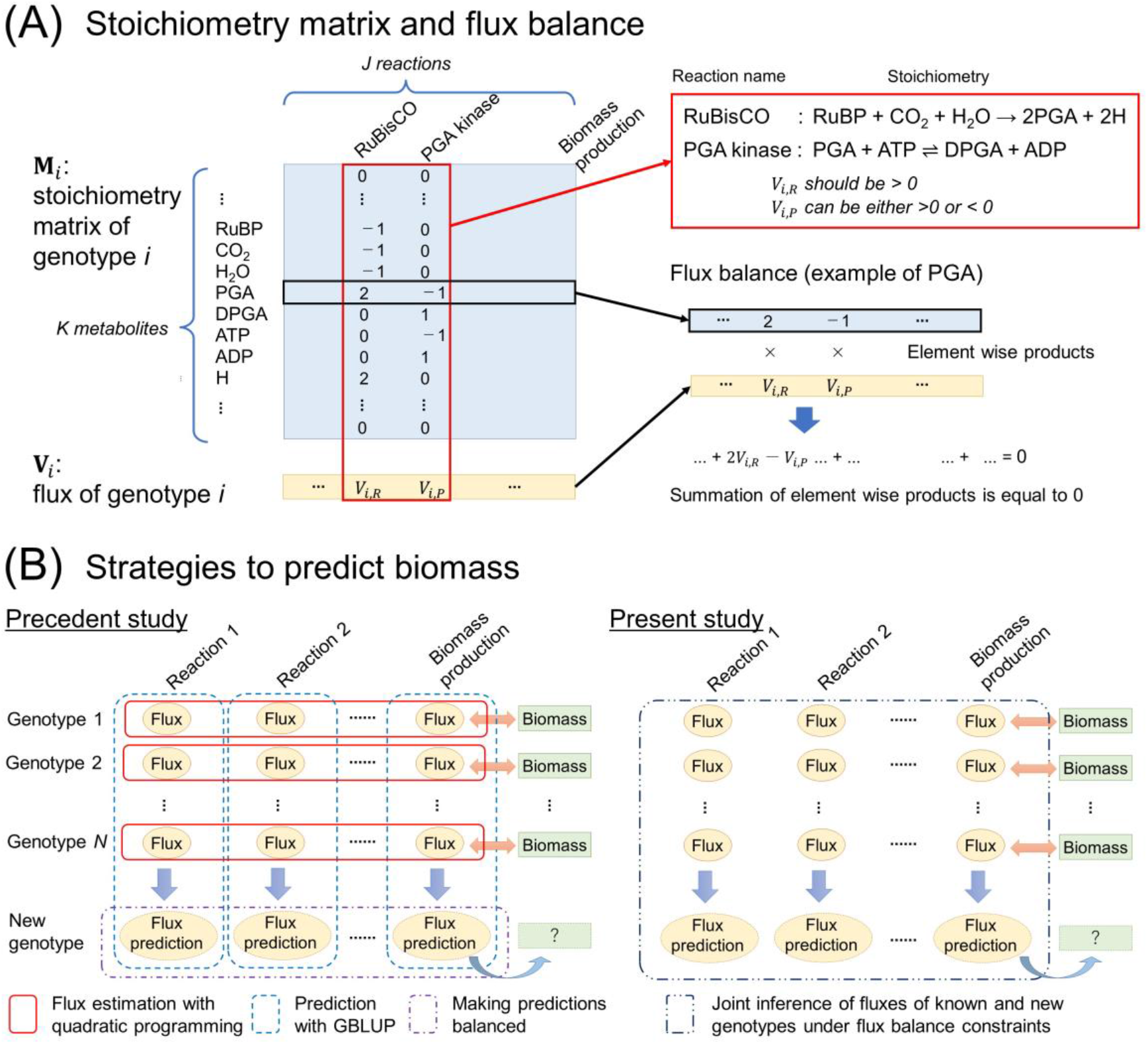
An overview of the proposed model and its analysis. (A) Diagram of the stoichiometry matrix and flux balance. Here, two reactions (RuBisCO and PGA kinase) and eight metabolites related to the reactions are shown. Flux balance analysis assumed that all the metabolites in the stoichiometry matrix are balanced. Note that biomass itself is not included in the stoichiometry matrix. (B) Strategy comparison between the previous study (Tong *et al*., 2020) and the current study. The precedent study strategy consists primarily of three steps: (1) Estimation of the flux of each genotype under flux balance constraints employing quadratic programming. (2) Prediction of fluxes of new genotypes employing genomic BLUP (GBLUP). (3) Making predicted fluxes balanced employing quadratic programming and predicting the biomass of new genotypes with the predicted flux of the biomass production. In contrast, the proposed model in the present research conducts these steps in one step employing a Bayesian hierarchical model.

Each metabolite’s production and consumption are balanced according to the flux balance analysis principle. This can be written as an equation such as

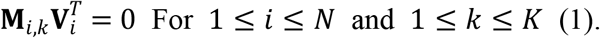

where **M**_*i,k*_ denotes the *k*_th_ row of **M**_*i*_. Eq. (1) is also illustrated in Fig. 1A. The flux of the *J*_th_ reaction (i.e., reaction for the target trait), *V_,J_*, is assumed to be proportional to the observed phenotypes of the target trait, **Y.** Here, **Y** is an *N*-length vector that contains the phenotypes of all genotypes.

To summarize, Tong *et al*., (2020) estimates **V**_*i*_ independently for each genotype so that Eq. (1) is filled and the ratio between *V_i,J_* and *V*_*Col*–0.*J*_, a flux of the reference genotype Columbia-0, is matched the ratio between *Y_i_* and *Y_Col-0_*. (Fig. 1B). The flux estimated in this manner is referred to as **Vt** in the following. See Section 6.2 for the explanation of prediction by Tong *et al*., (2020).

In the present study, **V**_*,J*_ is connected with **Y** as

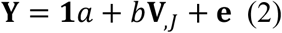

where **1** indicates the vector of 1 and **e** is a gaussian noise, that is, 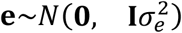 where **0** indicates the vector of 0 and **I** is an identity matrix.

The purpose of the proposed model is to estimate **V**_*,J*_ under the constraint of Eq. (1). To treat the constraint in a Bayesian hierarchical model, Eq. (1) is rewritten as

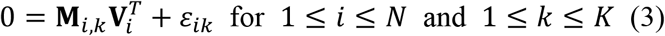

where 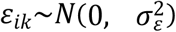. Eqs. (2) and (3) constitute the likelihood of the model. The prior distributions of 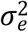 and 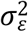 are half-Cauchy distributions written as,

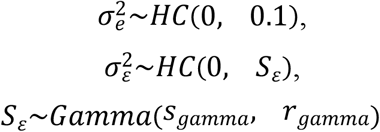

respectively. Here, *s_gamma_* and *r_gamma_* denote the shape and rate parameters set to 0.1 and 1.0, respectively.

Positive values of flux (**V**) mean that the reaction flows from the left to the right hand, whereas negative values mean the opposite direction. Some reactions can occur in both directions, whereas some only occur in either direction (Fig. 1A). Let *J_n_* denote the set of reactions that can proceed in both directions and *J_r_* denote the set of reactions that have a positive flux restriction. Then, the prior distributions of *V* are written as,

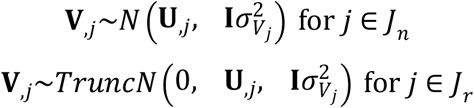

where **U**_*,j*_ defines the genetic component of **V**_*,J*_., and *TruncN* denotes truncated normal distributions. The prior distribution of **U**_*,j*_ is written as

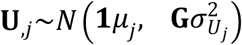

where *μ_j_* is a known mean value described later and **G** is a genomic relationship matrix that is defined with single nucleotide polymorphisms (SNPs).

The unobserved phenotype *Y_i_* of new genotype *i* can be predicted as

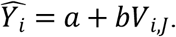

Here, *V_i,J_* can be estimated from the constraints of Eq. (3) even when the phenotype *Y_i_* is unavailable, and from *U_i,J_* which can be predicted with **G**. In Fig. 1B, the outline of the proposed model is illustrated with the precedent approach by Tong *et al*., (2020).

Based on prior physiological knowledge, some fluxes may be constrained by other fluxes. For example, in the biomass prediction proposed by Tong *et al*., (2020), it was supposed that

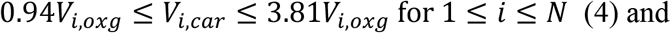

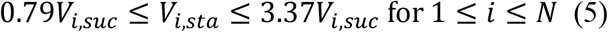

where *V_i,oxg_, V_i,car_, V_i,suc_*, and *V_i,sta_* denote the fluxes of oxygenation, carboxylation, sucrose synthesis, and starch synthesis reactions, respectively. Such constraints are referred to as “additional constraints” in the following. However, in the case of Arabidopsis biomass, these additional constraints resulted in erroneous predictions. This issue is mentioned in the Results and Discussions

## 3. Algorithm and implementation

All statistical inferences were conducted with R (ver. 4.1.3 on Windows machines or ver. 4.2.2 on Ubuntu 18.4 machines) (R Core Team, 2022). The posterior distributions of the proposed models were calculated using Hamiltonian Monte Carlo, which was implemented by rstan (ver. 2.21.5 or 2.21.7). The number of iterations was 5000, the length of the warm-up was 4000, and the thinning was 1. NUTS (No-U-Turn Sampler) was used for sampling. Because both **V** and **U** are high-dimensional (*N* by *J* matrices), to facilitate exploration of the posterior distribution, two modifications were made from the base model defined above when implementing the model using stan. These modifications are illustrated in subsections 3.1 and 3.2.

### 3.1. Scale and location

The scales 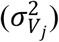 and locations (*μ_j_*) of flux are considerably different among reactions, which can hamper the efficient exploration of posterior distributions. Thus, using prior knowledge, the scales and locations were changed to be comparable across reactions. Suppose that rough estimates for 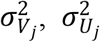, and *μ_j_*, 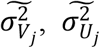, and 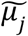, can be obtained from the literature and/or preceding experiments. Define *δ_j_* as

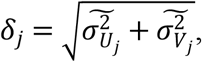

then divide **V** and **U** with *δ_j_* as **V***_,j_′* = **V**_*,j*_/*δ_j_* and **U***_,J_′* = **U**_*,j*_/*δ_j_* yielding prior distributions of **V***_,j_′* and **U***_,j_′* represented as

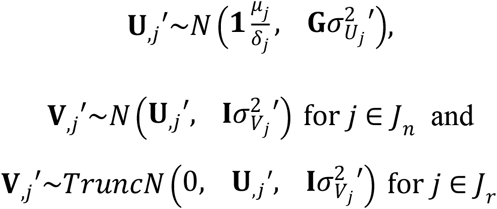

where 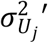 and 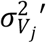 correspond with 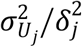 and 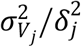, respectively. The location of **V**_*,j*_′ was then arbitrarily adjusted to 10 using 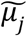 as

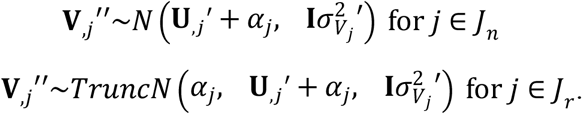

where 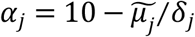.

Inference was conducted using **V***_,j_″* and **U***_,j_′*. The constraint of Eq. (1) becomes

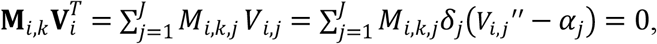

that is,

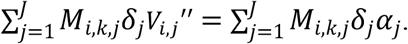

Thus, Eq. (3) can be written as

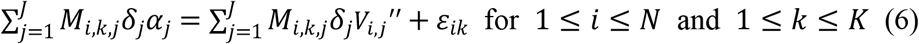

where 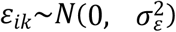. Eq. (2) is rewritten using **V***_,J_″* as

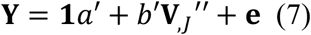

The prior distributions of *a’, b’*, and variances are half-Cauchy distributions written as,

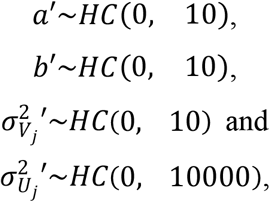

respectively.

### 3.2. Weight in the likelihood

Eqs. (6) and (7), terms for observations and constraints, respectively, define the likelihood of the model. However, the difference in dimensions (the observations consist of *N* components while the constraints consist of *N* × *K* components) can cause underfitting to Y if these two terms are combined naively in the log likelihood function. Thus, a parameter to control the relative weight between the observations and constraints was introduced to the log likelihood:

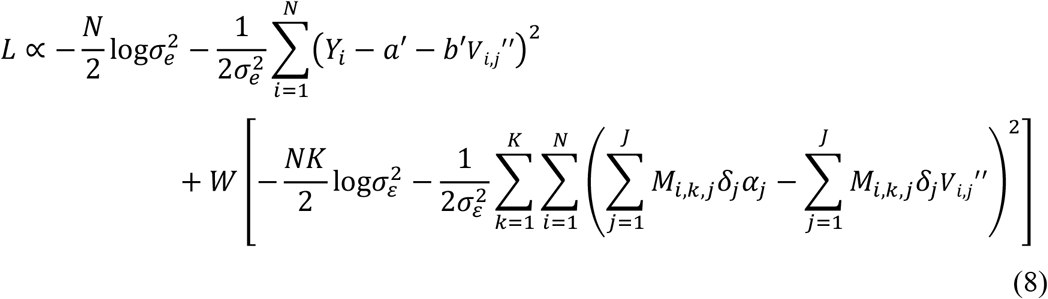

where *W* is the weight parameter. The effect of *W* was investigated using simulated and real data.

## 4. Real data analysis

The real data used in this study were the same as the data used by Tong *et al*., (2020). The target trait is the biomass of *Arabidopsis thaliana* evaluated under the 12 h-light optimal nitrogen conditions (Sulpice *et al*., 2013). The number of genotypes (*N*) was 67, including Columbia-0. Arnold and Nikoloski (2014) created the stoichiometry matrix (**M**) of the reference genotype relevant to biomass. The matrix originally contained 407 metabolites and 549 reactions. A stoichiometry matrix with 350 metabolites and 336 reactions (i.e., K = 350 and J = 336) was obtained by eliminating reactions with zero flux. The matrix was sparse; the medians of the number of metabolites per reaction and the number of reactions per metabolite were 4 and 2, respectively. The *J*_th_ column of the stoichiometry matrix contains genotype-specific coefficients, and Tong *et al*. (2020) estimated the coefficients for the genotypes other than the reference. Horton *et al*. (2012) genotyped the 67 genotypes’ genome-wide SNPs. The genomic relationship matrix **G** was calculated from exome SNPs by Tong *et al*. (2020). All real data including **G** are available at https://github.com/Hao-Tong/netGS.

The prediction ability of the proposed model was assessed with 3-fold cross-validation (CV). The genotype divisions were done following Tong *et al*., (2020). This precedent study conducted a 3-fold CV 50 times and opened the genotype divisions of all CVs at the above-mentioned site. Following the first 20 CVs of Tong *et al*. (2020), a 3-fold CV was carried out 20 times in the current study. R package snow (ver. 0.4-4) was used to parallelize CVs.

Modifying scales and locations as described in subsection 3.1 requires rough estimates for 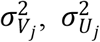, and *μ_j_* (i.e., 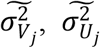, and 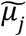). To obtain these estimates, the following linear model was fitted to the training data:

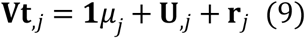

where **Vt**_*,j*_ is the fluxes of reaction *j* estimated by Tong *et al*., (2020), 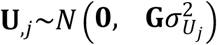, and **r**_*j*_ is the residual following 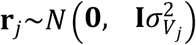. Note that **Vt**_*,j*_ was estimated using quadratic programming for each genotype without information on SNP genotypes. It is also noteworthy that in this real data analysis, **Vt**_*,j*_ only took into account the genotype fluxes from the training data and did not use those from the testing data for this estimation. The solutions of 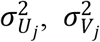, and *μ_j_* were obtained employing an R package rrBLUP ver. 4.6.1 (Endelman JB, 2011).

The unobserved phenotype *Y_i_* of new genotype *i* was predicted as

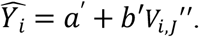

Prediction accuracy was investigated using the Pearson correlation coefficient (R) between *Y_i_* and 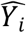. Following Tong *et al*., (2020), R was calculated for each fold of each CV As the weight parameter *W*, six values (0.16, 0.32, 0.48, 0.64, 0.80, and 0.96) were compared. Differences in accuracy between methods were tested with the Mann–Whitney–Wilcoxon test using the R function wilcox. test.

## 5. Simulation analysis

To assess the prediction ability of the proposed model, data were simulated by mimicking the real data. First, employing the model of Eq. (9), the location (*μ_j_*) and variance components (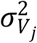 and 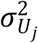) were estimated from all genotypes (*N* = 67). Then, **U**_*,j*_ and **V**_*,j*_ were simulated from

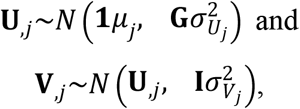

respectively. If the flux of reaction *j* is restricted to be positive, samples were drawn until the restriction was met. **M** was the stoichiometry matrix of the reference genotype (Columbia-0). However, for each genotype, the last non-zero reaction of each metabolite was modified as follows to fill the constraints of Eq. (1):

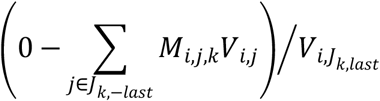

where *J_k,–last_* indicates the set of reactions in **M** that are non-zero for metabolite *k* except for the last reaction, *J_k,last_*. For example, when the metabolite *k* has non-zero coefficients in **M** at reactions 4, 18, and 125,

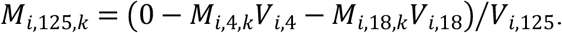

The observed phenotypes (**Y**) were then generated by

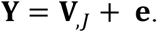

Herein, **e** was drawn from the normal distribution. The variance of the normal distribution was set to 25% of that of **V**_*,J*_.

To facilitate model fitting in simulations, a small noise, *ψ_ik_*, was applied to the left hand of Eq. (6). That is,

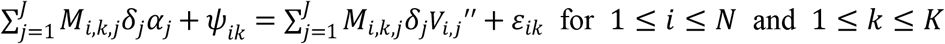

where *ψ_ik_*~*N*(0, 0.0001) of which the variance was determined arbitrarily.

With this process, 10 simulated data sets were generated. Then, a 3-fold CV was carried out for each data set. Genotype divisions of CVs followed the first 10 CVs of Tong *et al*., (2020). As performed in the real data analysis, prediction accuracy was evaluated using R, which was calculated for each fold of each CV. Six values (0.16, 0.32, 0.48, 0.64, 0.80, and 0.96) were compared as the weight parameter *W*. Differences in accuracy between methods were tested with the Mann–Whitney–Wilcoxon test.

## 6. Methods compared

### 6.1. Genomic BLUP (GBLUP)

Genomic BLUP (GBLUP), the most widely employed method in genomic prediction, was carried out employing the observed phenotypes (**Y**) as the response variable. The model can be written as

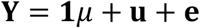

where 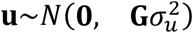 and 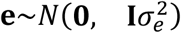, respectively. No information about metabolites and metabolic networks was used. The rrBLUP package ver. 4.6.1 was used to execute.

### 6.2. Quadratic programming (QP)

This method was proposed by Tong *et al*., (2020). First, using the model Eq. (9), GBLUP predicts fluxes of genotypes in testing data (see also Figure 1B). The predicted fluxes, *U_test,j_*, however, do not fill the constraints of Eq. (1). Thus, the steady state fluxes, *V_test,j_*, are estimated using QP in the R package CVXR (ver. 1.0-10) (Figure 1B). The objective function and constraints of this optimization are

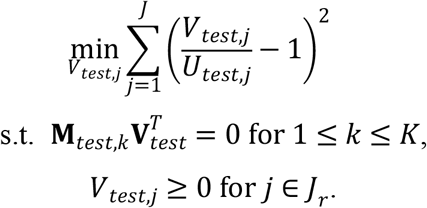

*V_test,J_* at the steady state was then treated as the predicted value for Y. Although the constraints of Eqs. (4) and (5) were applied in the original method proposed by Tong *et al*., (2020), here QP was performed without these additional constraints due to the reasons described in the outcome and Discussion.

### 6.3. MegaLMM

MegaLMM is a multi-trait mixed model that is based on the factorization of genetic covariances between traits (Runcie *et al*., 2021). This method treated estimated fluxes (**Vt**_*,j*_) and **Y** as traits, and the following multi-trait mixed model was fitted:

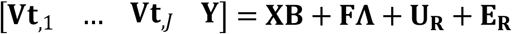

where

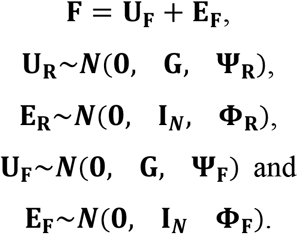

The second and third arguments in the normal distribution *N* denote the covariances between rows and columns, respectively. **X** represents the design matrix, which only included the intercepts for each trait, and **B** represents the fixed effects. **F** is an *N* x *P* latent factor matrix, and **Λ** is a *P* × (*J* + 1) factor loading matrix. **Ψ_R_** and **Φ_R_** are diagonal matrices of dimension *J* + 1 and **Ψ_F_** and **Φ_F_** are diagonal matrices of dimension *P*. Here *P* is the dimension of the latent variables. The prior distributions of parameters followed the vignette of the R package MegaLMM, which can be found at (https://github.com/deruncie/MegaLMM/blob/master/vignettes/Running_MegaLMM.Rmd). **Y** was scalded before fitting. The total number of iterations was 10,000, thinning was 2, and the posterior inference was performed on the last 500 samples. *P* was set to 50, 80, or 160. The R package MegaLMM (ver. 0.1.0) was used to fit the model.

## 7. Results and discussion

The outcome of CVs employing simulated data is presented in Fig.2. QP and the proposed models generally showed greater accuracy than GBLUP, suggesting that the information of metabolic networks is advantageous in predicting the target trait phenotype. The accuracy of the proposed model depended on the weight parameter *W*. The accuracy showed a convex curve peaking at *W* = 0.64. The proposed models were able to predict more accurately than QP at *W* = 0.48 and 0.64, suggesting that the proposed models potentially have superior prediction ability than QP as intended. MegaLMM was less accurate than GBLUP, although MegaLMM also used the multivariate information of fluxes. Perhaps, the factorization of genetic covariance has no meaning in these data and might make inference unstable because genetic covariances among fluxes and the target trait were not simulated.

**Fig. 2.**
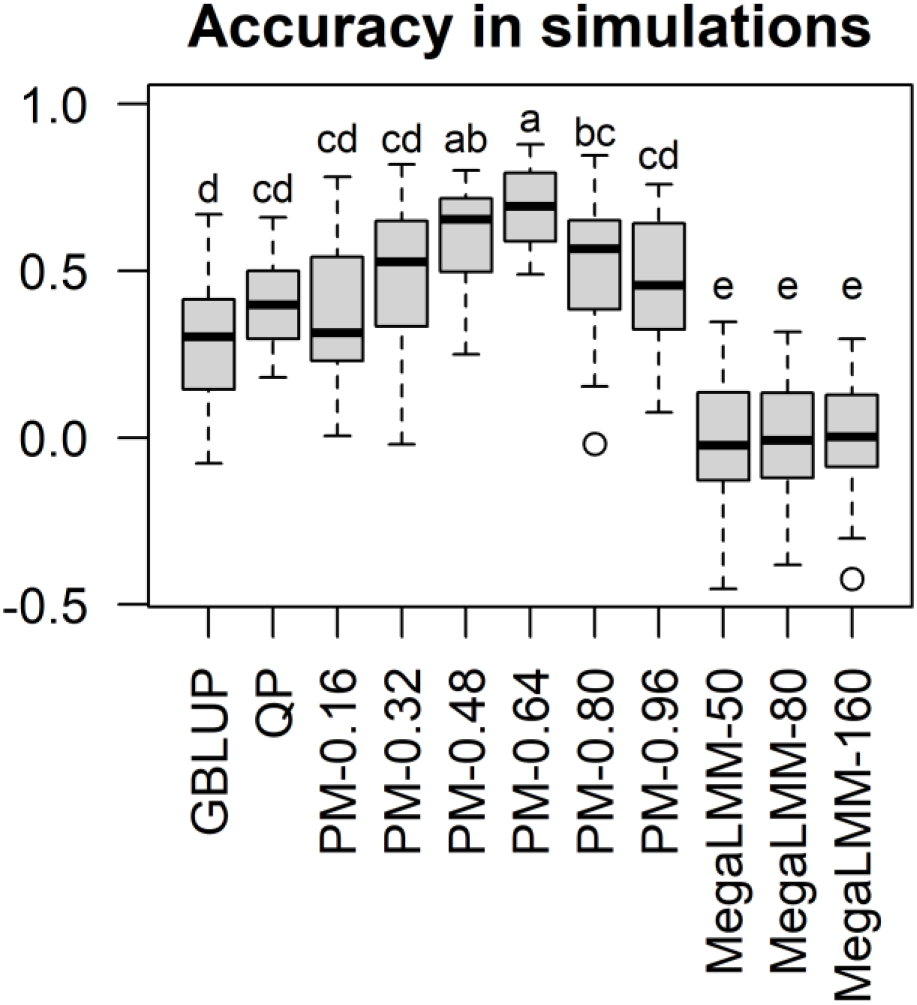
Accuracy in cross-validations employing simulated data. Pearson correlation coefficients between the predicted and observed phenotypes are presented. There are significant differences between different characters (P < 0.05, after Bonferroni correction). GBLUP, genomic BLUP; QP, quadratic programming; PM-*X*, proposed model with the weight parameter *W* = *X*; MegaLMM-*X*, MegaLMM with the number of latent factors = *X*.

The results of CVs employing the real data are presented in Fig. 3. Here, the QP was significantly inferior to GBLUP. The medians of the accuracies were 0.012 and 0.239, respectively. The accuracy of the proposed model showed a convex curve with *W* as observed in the simulations, but the peak shifted to *W* = 0.32. At this *W* value, the proposed model performed significantly better than GBLUP (the median 0.367 vs. 0.239). MegaLMM was equivalent to GBLUP in terms of accuracy. Fig. 4 illustrates how strictly the constraint of Eq. (6) was filled. The Pearson correlation between 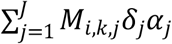 and 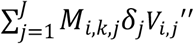 was almost 1.0 at each *W* value, suggesting that the fluxes were at steady state.

**Fig. 3.**
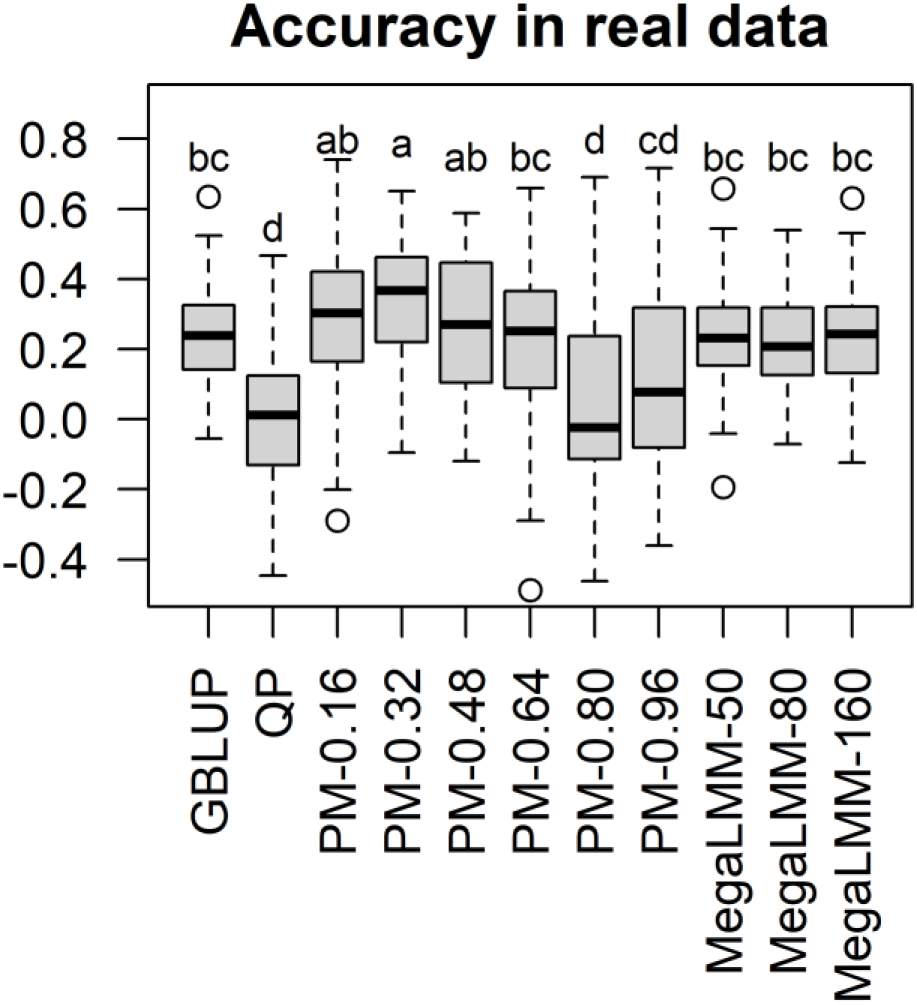
Accuracy in cross-validation employing real data. Pearson correlation coefficients between the predicted and observed phenotypes. There are significant differences between different characters (P < 0.05, after Bonferroni correction). GBLUP, genomic BLUP; QP, quadratic programming; PM-*X*, proposed model with the weight parameter *W* = *X;* MegaLMM-*X*, MegaLMM with the number of latent factors = *X*.

**Fig. 4.**
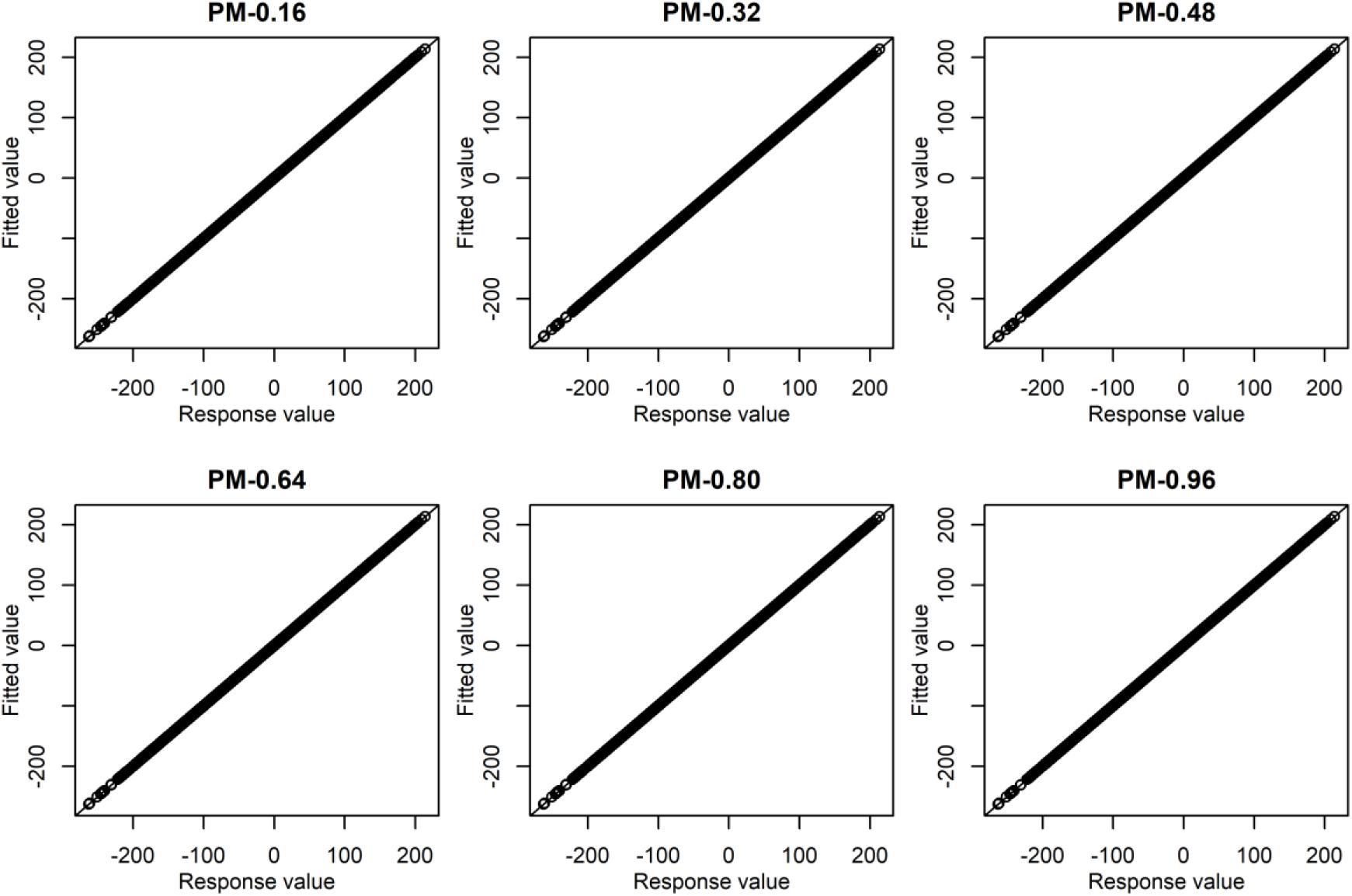
Fitting status in the constraint term. The x-axis is 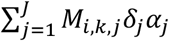 and y-axis is 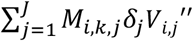 of Eq. (6). PM-*X* denotes the proposed model with the weight parameter *W* = *X*. Results from all CVs were pooled. The diagonal line is the 1:1 line.

When comparing the real data results to the simulation results, two major differences were observed: (1) Fewer advantages in using flux balance information and (2) shift of optimal *W* values. The former may arise because of inadequate or inaccurate information on the metabolic networks of biomass production of Arabidopsis. That is, not all metabolites and reactions may have been listed, and/or some stoichiometry coefficients may have been calculated incorrectly. It will be a question for future studies as up to what extent such deficiencies in network information affect prediction accuracy. Regarding the second point, the relative importance between the observation (Y) and constraints (flux balance) may differ between simulated and real data because the weight parameter *W* controls the relative contributions of these two terms to likelihood, as shown in Eq. (8). Intuitively, if the observations are more reliable as an information source to infer fluxes (**V**) than the constraints, lower *W* values may be preferable and if the observations are less reliable, higher *W* may be preferable. Supposing that the reliability as an information source could be quantified with heritability, the outcome may suggest that the heritabilities of fluxes in real data were lesser than those employed in the simulations. Although the heritabilities in the simulations were taken from those estimated from real data, the estimates must have been overestimated due to the small data size, and overestimation for the fluxes would be more influential on inference than overestimation for the observations (Y) due to the difference in dimensions (*J* vs. 1).

The unexpectedly low accuracy of QP in the real data analyses was drastically advanced by introducing additional constraints illustrated in Eqs (4) and (5). The median accuracy was 0.414, which was siginificantly greater than that of GBLUP and the proposed models. However, it was found that the predicted values hardly fluctuated among CVs. The mean of the Pearson correlations between predicted values obtained in different CVs was 0.999, indicating that the fluctuations in prediction accuracy reported by Tong *et al*., (2020) were caused by genotype division variation in CVs. Moreover, even when the genomic relationship matrix (**G**) was permutated, the prediction accuracy was still high (median 0.424), suggesting that the prediction accuracy was achieved without using genomic information. Further research confirmed that the constraint of Eq. (4) is enough to cause these phenomena (data not shown). Although it is unclear why this constraint causes the phenomenon, these findings suggest that when using flux balance for prediction, it is important to be cautious about putting constraints on fluxes.

Both the simulated and real data analyses revealed that the proposed model is capable of improving the prediction accuracy of quantitative traits. However, there are three concerns about the proposed model to be mentioned. The first is the computational cost. Hamiltonian Monte Carlo simulations typically took 21 h for model fitting (Windows 11 machine with Core (TM) i7-12700K 3.61 GHz), which is significantly slower than GBLUP (took <1 sec.). The high computational cost of the proposed model is not surprising because the model includes *J* (336) random effects of which each dimension is *N* (67) (**U**). The second complication is how to determine *W* before analysis. A grid search using CV could be a viable solution, albeit at a higher computational cost. Another solution will be to put a prior distribution on *W* and infer from the data. The last problem is the assumption of independency among the fluxes and target traits. Because fluxes would be correlated with each other, ignoring this dependency can make inference unstable. The factorization adopted in MegaLMM will be useful to mitigate this issue. Such improvement in models will be needed to make the proposed models more applicable to various traits and species.

In summary, the proposed model was shown to be capable of improving the prediction accuracy using metabolic network information without employing additional constraints on fluxes. Although there are still problems to be addressed, this research supports the idea of Tong *et al*., (2020) that metabolic network information can be used to predict phenotypes of quantitative traits.

## Acknowledgments

The author thanks Ms. Yuuri Onogi for her support in conceiving and conducting this study.

## Data and source code availability

All real data are available at https://github.com/Hao-Tong/netGS. Stan and R scripts required to reproduce this study are available at https://github.com/Onogi/MetabolicModeling.

